# Back to the Meadow Brown: eyespot variation and field temperature in a classic butterfly polymorphism

**DOI:** 10.1101/2022.07.08.499313

**Authors:** Sophie Mowbray, Jonathan Bennie, Marcus W. Rhodes, David A.S. Smith, Richard H. ffrench-Constant

## Abstract

Since the classic work of E.B. Ford, alternate hypotheses have focused on explaining eyespot variation in the Meadow Brown butterfly strictly as a genetic polymorphism and the role of temperature in this classic example of natural selection has therefore been overlooked. Here we use large and continuous field collections from three sites in the UK to examine the effect of field temperature on total eyespot variation using the same presence/absence scoring as Ford. We show that higher developmental temperatures in the field lead to the disappearance of the spots visible while the butterfly is at rest, explaining Ford’s original observation that hindwing spotting declines across the season as temperatures increase. Analysis of wing damage supports the historical hypothesis that hindwing spots confuse aerial predators. However, as hindwing spotting declines over the season, a ‘trade-off’ is suggested between their role in deflecting predators early in the season and their later developmental cost. In contrast, the large forewing eyespot is always present, scales with forewing length and its variation is best explained by day of the year rather than developmental temperature. As this large forewing spot is thought to be involved in ‘startling’ predators, its constant presence is therefore likely required for defence. We model annual total spot variation with phenological data from the UK and derive predictions as to how spot patterns will continue to change under increasing summer temperatures, predicting that spotting will continue to decrease both across a single season and year or year as our climate warms.

**Summary statement:** We show that a long-held example of ‘genetic’ polymorphism, eyespot variation is the Meadow Brown butterfly, is correlated with field temperature during butterfly development.

## Introduction

Phenotypic variation is driven by a balance between genetic and environmental factors (Pfennig et al., 2010; Schneider and Meyer, 2017). In their pioneering studies, E.B. Ford, W.H. Dowdeswell and R.A. Fisher used eyespot variation in the Meadow Brown butterfly, *Maniola jurtina*, to help define the term ‘genetic polymorphism’ and then went on to establish the nascent field of population genetics (Dowdeswell et al., 1949; Dowdeswell and Ford, 1955; Dowdeswell et al., 1957; Dowdeswell et al., 1960; Dowdeswell and McWhirter, 1967). Most of these field studies assumed that both within (intra-) and between (inter-) season eyespot variability in this butterfly was associated with genetic variation (Brakefield and van Noordwijk, 1985; Creed et al., 1964; Dowdeswell, 1981). In other words that the butterfly existed as a series of high and low spot ‘morphs’ with spotting being under genetic control as a classic example of a genetic polymorphism. Several hypotheses were subsequently proposed to explain the anticipated differential survival of these high and low spotted morphs, including differential predation or infection (Dowdeswell, 1981) and differential rates of morph development driven by pleiotropic effects of the wing patterning genes involved (Beldade et al., 2002; Brakefield and French, 1995; Brakefield and Shreeve, 1992; Connahs et al., 2019; Dhungel et al., 2016; French and Brakefield, 1995; Iwasaki et al., 2017; Koch et al., 2003; Monteiro et al., 1997a; Monteiro et al., 1997b; Monteiro et al., 2013; Otaki, 2020; Reed and Serfas, 2004; Sekimura et al., 2015; Zhang and Reed, 2016). However, some studies also hinted that the environment might also be important in controlling variation. First, measurements of heritability not only differed between the sexes but were also higher at higher temperatures (Brakefield and van Noordwijk, 1985). Secondly, local environmental changes such as the cessation of grazing (Dowdeswell and Ford, 1955; Dowdeswell et al., 1957) or prolonged drought (Bengston, 1978; Dowdeswell et al., 1960) also sometimes appeared to unexpectedly change spot patterns, perhaps due to changes in the associated micro- or macro-climate. In terms of likely environmental effects, ‘phenotypic plasticity’ can be defined as the ability of individual genotypes to produce different phenotypes when exposed to different environmental conditions, one of which is temperature (Pfennig et al., 2010; Schneider and Meyer, 2017). In insects, this plasticity is well documented in the laboratory with increases in temperature either enhancing (Brakefield et al., 1998) or diminishing (Zhang et al., 2020) spot-like pattern elements. For butterflies, the critical period for determining the development of wing pattern formation is early pupation (Brakefield et al., 1998). However, most studies of eyespot variation in butterflies come from temperature-controlled laboratory experiments (Brakefield et al., 1996; Brakefield et al., 1998) and the effects of *field* temperature on phenotypic variation therefore remain unknown. Here we examine the simple hypothesis that field temperatures during pupation control eyespot variation (defined by spot presence/absence) in the Meadow Brown butterfly. To test this, we re-examine the exact presence/absence type of data (spot scoring) collected by Ford and others, to examine the role of field temperature in explaining their historical and formative observations on spot variation both within and between seasons.

As well as repeating the presence/absence scoring of Ford, we also wanted to acknowledge the likely role of wing length in determining the size of the large and omni-present spot on the forewing of both sexes (here termed the large ‘compound’ spot 2/3), thought to be involved in predator deterrence. However, frustratingly, the substantial volume of work by E.B. Ford and others was conducted in a manner which makes their studies not only hard to reproduce or re-examine but also difficult to compare with the subsequent work of others. For example, Ford often collected butterflies with a group of observers in the field, with different observers scoring the butterflies directly in the net and then releasing them(Dowdeswell, 1981; Dowdeswell et al., 1949; Dowdeswell and Ford, 1952; Dowdeswell and Ford, 1955; Dowdeswell et al., 1957; Dowdeswell et al., 1960; Dowdeswell and McWhirter, 1967). This means not only that spot presence/absence was scored by multiple observers on living animals but also that we cannot return to many of their historical specimens to re-score them as they were not captured and set. They also only scored relatively small numbers of female butterflies over different dates at different sites and ignored males altogether. Finally, the numbering used for spot scoring differs between their studies and those of other subsequent authors, making direct comparisons extremely confusing, with most studies (including Ford’s) only examining the presence/absence of spots on the female hindwing and ignoring forewing spotting entirely.

To get around these problems and to increase reproducibility here we made several simple adjustments to their spot presence/absence scoring. First, we collected large numbers of insects day on day throughout the entire flight period and then ‘side-set’ all butterflies of both sexes, providing a permanent resource to which we and others can return. Second, all butterflies were scored by the same observer, removing the variation shown to be introduced via scoring with multiple observers (Brakefield and Dowdeswell, 1985). Third, we adopted a strict definition of ‘presence’ as raised eyespot scales visible under a 10x hand-lens on a set butterfly, something not readily achievable in the field with a struggling butterfly in a net and thus giving a strict and reproducible definition of presence/absence. Fourth, we simplified the numbering of all ten spots with five candidate spot positions scored on the forewing (spots one to five) and five on the hindwing (spots six to ten). Fifth, we scored all ten spot positions in both sexes in very large and continuously collected populations from a range of UK locations over several years, both historical (1980s-1990s) and current (2020). Finally, as well as scoring the presence/absence of all ten spots we also examined the likely role of wing length in determining the size of the larger compound spot termed 2/3, which is invariably present on the forewing of both sexes and therefore cannot be scored as present/absent. Using this revised scoring method, here we show that field temperature 35 days prior to butterfly capture (here termed ‘developmental temperature’ and assumed to be the temperature at the time of pupation) explains both why the small spots visible at rest decline if frequency over the season and within hotter summers. We use this temperature-based model to predict how spot frequencies are likely to change in the UK with our warming climate.

## Materials and methods

We wanted to repeat the scoring method of Ford to look at the effect of field temperature on the probability that any one of the ten different eyespots are present/absent on the wing of both sexes of field collected Meadow Browns. To this end we collected large and continuous (daily collections) of butterflies from three sites in the UK over several years, both historical and present day.

### Field collections and spot scoring

Butterflies of the same sub-species of *M. jurtina insularis* (Thomson, 1969) were collected from three different sites across the UK. 1) Eton in Berkshire (grid reference SU965775 with butterflies collected every year from 1988-1993). 2) Buckingham in Buckinghamshire (grid reference SP687339 from 1988-1991). 3) Chycoose (near Truro) in Cornwall (grid reference SW804390 and year collected 2020). We then ‘side-set’ these large collections (series) of both male and female butterflies with only one underside of each specimen visible from above. Side-setting means we can readily score all spot positions on each individual whilst reducing the very large space necessary to store all specimens in a museum for further reference. To reduce the variation shown to be introduced by multiple observers scoring the same specimens by eye, and the problems of scoring in in the field (see above), we adopted the following improvements to Ford’s original scoring protocol. 1) We numbered all spot positions from one to ten (Supplemental Fig. S1). With spot positions numbered 1-5 on the forewing and positions 6-10 on the hindwing. Spots 2/3 on the underside of the forewing are usually joined and were therefore scored as a ‘compound’ spot (see below). We therefore looked at *all* spots on both the fore- and hindwing and not just the hindwing spots scored by Ford and many other authors (Dowdeswell, 1981; Dowdeswell et al., 1949; Dowdeswell and Ford, 1952; Dowdeswell and Ford, 1955; Dowdeswell et al., 1957; Dowdeswell et al., 1960; Dowdeswell and McWhirter, 1967). 2) All scoring was performed by a single observer (D.A.S) defining spot ‘presence’ as the presence of at least one melanic raised scale visible under a 10x hand lens, at the predicted spot location, and the lack of any such raised scale as ‘absence’. This ensures a *precise and repeatable* method of scoring spot presence/absence not possible in the field with living animals in a butterfly net. Using a single observer to derive the entire data set in the current study therefore removes any possibility of variation introduced by different scorers (Brakefield and Dowdeswell, 1985). 3) We scored all ten spot positions in both sexes (males and females) in all series. 4) We recorded the presence of wing damage on each of the four wings and whether it was ‘symmetrical’ (cutting through both sets of wings in the same pattern) or ‘asymmetrical’ (damage only confined to one set of wings or wing). We also measured the forewing length (in mm using Vernier callipers) of the large and continuous (with daily collection) series of Cornish females collected from Chycoose Farm, Truro, in the summer of 2020, alongside both the width and height of the compound spot 2/3. Width (*a*) and height (*b*) were then used to estimate the total area (*A*) of spot 2/3 using the formula *A* = *π* a b.

### Wing damage scoring and analysis

To re-examine the hypothesis that eyespots deflect predator attack we also scored wing damage. Wing damage (fore- or hindwing) was scored for all butterflies collected from the single site Eton, across all years. Damage (‘beak’ marks or nicks to each wing) was scored as being absent (zero) or present (one), regardless of the level of damage present (e.g. number of nicks to each wing). Damage was also scored as symmetrical (passing through two or more wings with the same pattern) or asymmetrical. Results where then analysed with a binomial regression with damage scored as zero or one. The data presented in Fig. 4 are the model coefficients for each spot term. These are negative for a negative relationship (more spots leading to *less* damage) or positive for a positive relationship (more spots leading to *more* damage).

### Field temperature data

To correlate field temperature with the date of collection we used data from several sources. Temperature data were downloaded from the UK Meteorological office (MET) office for each of the locations and years when specimens were collected, during the flight season between 1st March and 31st October. For Buckingham and Eton, HadUK-Grid data was downloaded from https://www.metoffice.gov.uk/research/climate/maps-and-data/data/haduk-grid/datasets. This data gives the maximum and minimum air temperatures, measured over 24 h; the average of these two temperatures was used in the analysis. For the Chycoose Farm, Truro, Cornwall location, at time of analysis, the 2020 temperature data was not yet available, so instead temperature data from the closest weather station, located approximately 18 km away, in Camborne, was used instead.

### Principle components analysis of spot variation between sexes

All data analysis was conducted using RStudio (version 1.3.1093). Principle components analysis (PCA) was conducted to identify patterns across the ten wing spots, and to reduce potentially complex patterns into a more restricted number of principle components. The first principle-component, when run with all the data, was found to be associated with differences between the sexes. As most of the variation in wing spot patterns was due to sex differences, all subsequent analyses use only the female data. This is a standard approach taken during historic studies, as females are the more variable sex (Dowdeswell and Ford, 1952). The PCA was then repeated using only the females. The first principle-component was associated with the wing spot total, so the decision was taken to use the wing spot total as a measure of overall variation in subsequent analyses.

### Intra-seasonal spot number variation

A general linear model (or GLM) was used to test whether the specimens showed intra-seasonal shifts in wing spot patterns. The model included the wing spot total as the response variable, and day of the year of specimen collection and location as explanatory variables. The significance of each of these explanatory variables was tested using likelihood ratio tests. The model was subsequently plotted as a scatter plot, with the model as a regression line to allow visualisation of results.

### Sliding window analysis to determine ‘developmental temperature’

In other species, it has been shown that wing phenotypes are determined during the late instar larvae and early pupation (Beldade and Monteiro, 2021). The specimens used in this study were all caught in the field so the exact timing of their pupation is of course unknown. Therefore, it was necessary to identify the number of days prior to day of capture (the lag time) that the temperature best explains wing spot patterns. Different developmental temperatures were used in modelling, with lag times ranging from 1 to 75 days, and standard deviations of between 1 and 10. GLMs with wing spot total as the response variable, and developmental temperature, day of the year and location as explanatory variables were run. The Akaike Information Criterion (AIC) values of these different models were compared, and the lag time from the model with the lowest AIC value was used to determine the developmental temperature for all subsequent analyses. Plots were created of lag time against the AIC values, with each standard deviation plotted as separate lines, to allow visualisation of the results.

### The effect of developmental temperature on wing spot total and the presence/absence of individual spots

A linear mixed effect model was used to test the effect of developmental temperature on wing spot total. The mixed effect model, using the “lme4” package in RStudio, had wing spot total as the response variable, developmental temperature, location and day of the year as fixed effects, and year as a random effect. Model simplification, using likelihood ratio tests, was conducted to identify the minimum adequate model explaining variation in wing spot totals. Generalised linear mixed effect models were also used to test the effect of developmental temperature on the presence or absence of each of the individual wing spots. Each wing spot was modelled separately, with wing spot presence or absence as the response variable, developmental temperature as a fixed effect, and year as a random effect. A binomial error structure was used, as wing spot presence or absence is a binary variable. Location and day of the year were not included as fixed effects, as the results of the prior analysis on wing spot total found them to be non-significant. Model simplification, using likelihood ratio tests, was again conducted to identify the minimum adequate model explaining variation in individual wing spot presence or absence. Sigmoidal plots were then produced for each of the ten wing spots, using the package “sjplot” in RStudio, which plots the fixed effects of the model.

### Predicting general spatial and temporal trends in wing spot patterns from UK BMS data

Transect data was obtained from the UK Butterfly Monitoring Scheme (BMS), which has established over 2,500 transects across the UK since 1976. Fixed transect routes are walked between April and September, and all butterflies within 2.5 metres either side of the transect line and 5 metres ahead of the surveyor are recorded. The data used from the BMS included the location of each transect, the date it was walked, and the total count for the number of *M. jurtina* observed. Due to inconsistencies in the longevity of different transects, and the frequency of sampling, the full data set was filtered. Twenty transects (Table S2) were selected which provided a good spatial covering of the UK. The selected transects were all established prior to 1981, and were walked regularly each year, which allows temporal patterns to be identified.

### Phenological differences across UK

To account for potential differences in phenology (e.g. butterfly flight period) across the UK, the BMS count and date data were used to test whether there is any evidence of phenological differences in *M. jurtina* flight periods in different locations across the UK. If *M. jurtina* are adjusting their phenology in different parts of the country, this will affect the developmental temperatures that butterflies are exposed to, and therefore will affect the expected spatial patterns of wing spot totals. The selected transects were divided into ‘North’, ‘Southwest’ and ‘Southeast’ based on their location. The location of these selected transects, and how they were divided, was plotted using QGIS (3.16.2), and can be seen in Figure 3a. Scatter plots with day of the year against count were plotted for these localities separately. This plotting allowed any visual differences in the length of the flight period, or the peak of the flight period, to be identified. To compare the phenologies of these localities, the 2.5%, 25%, 50%, 75% and 97.5% quantiles of the day in the year were calculated, to see if the spread of the data differed. Each day of the year value was repeated by the count value, so that there was effectively a separate day of the year data point for every individual *M. jurtina* observed, to account for differences in population sizes. The difference between the 2.5% and 97.5% quantile values shows the spread of 95% of the day of the year data and was compared to look at differences in the length of the flight season. These quantile values were plotted as boxplots overlaying the scatter plots.

**Fig. 1.**
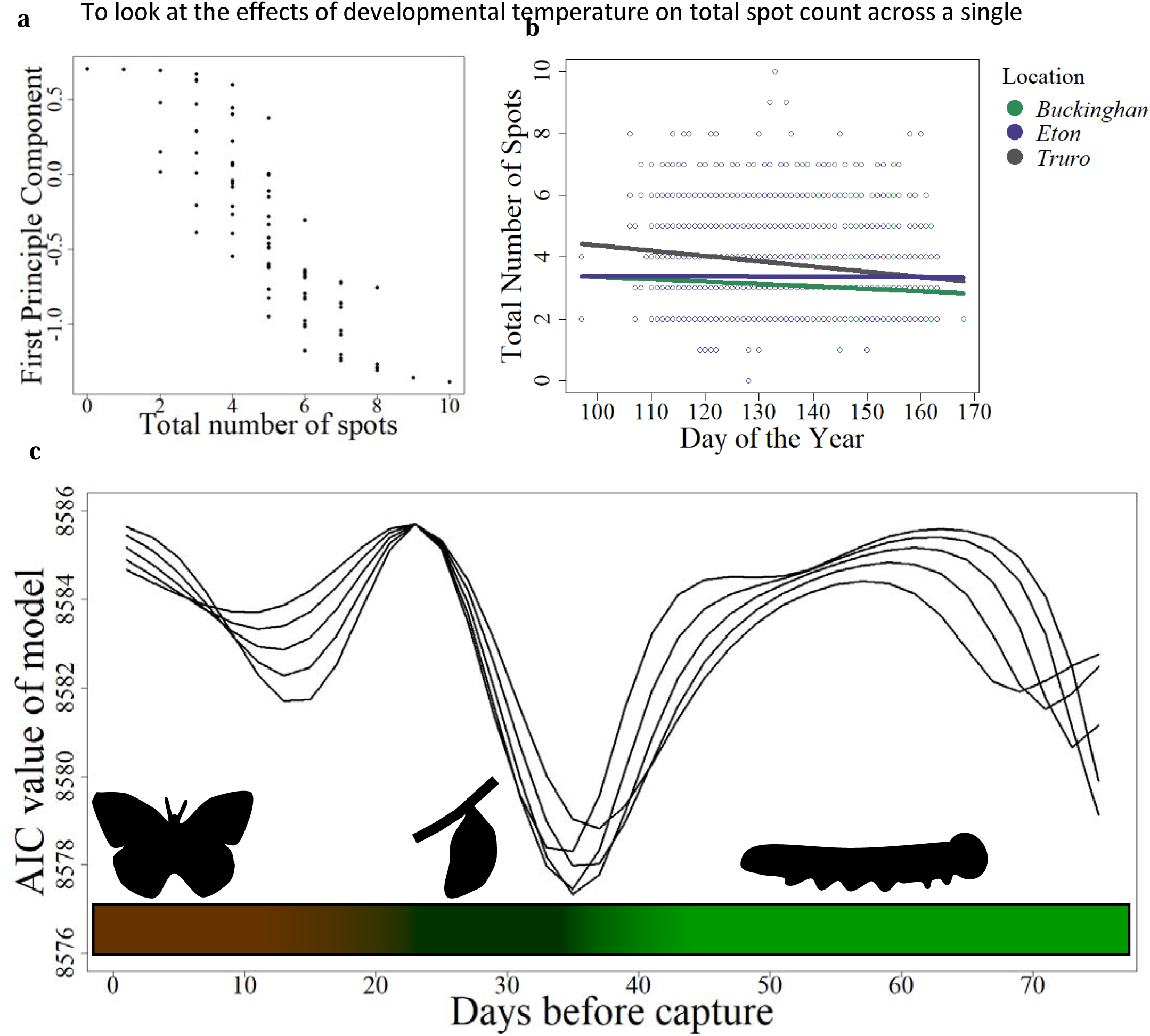
Temperature at early pupation correlates with the decline in female spottiness within a season. **a**, Principle components analysis shows that the first principle component is correlated with total number of spots. **b**, The total number of female spots declines through the season at all three sites (Eton, Buckingham and Truro). Day of the year is the number of days since the start of the flight season on 1 March. **c**, The Akaike information criterion (AIC) results from the models run lag times (days before specimen capture) of between 1 and 75 days, and standard deviations between 6 and 10 (shown as separate lines). The model was run with total number of spots as the response variable, and location, day of the year and developmental temperature, at the given lag and standard deviation, as the explanatory variables. The model with the lowest AIC value had a lag of 35 days with a standard deviation of 8. The coloured bar at the bottom of the plot reflects the likely life stages of the butterfly at each of the lag times, with brown showing the adult stage, dark green showing the pupal stage (∼28 days) and light green showing the larval stage. Data for female butterflies only, from all sites combined (n=2,504 butterflies).

**Fig. 2.**
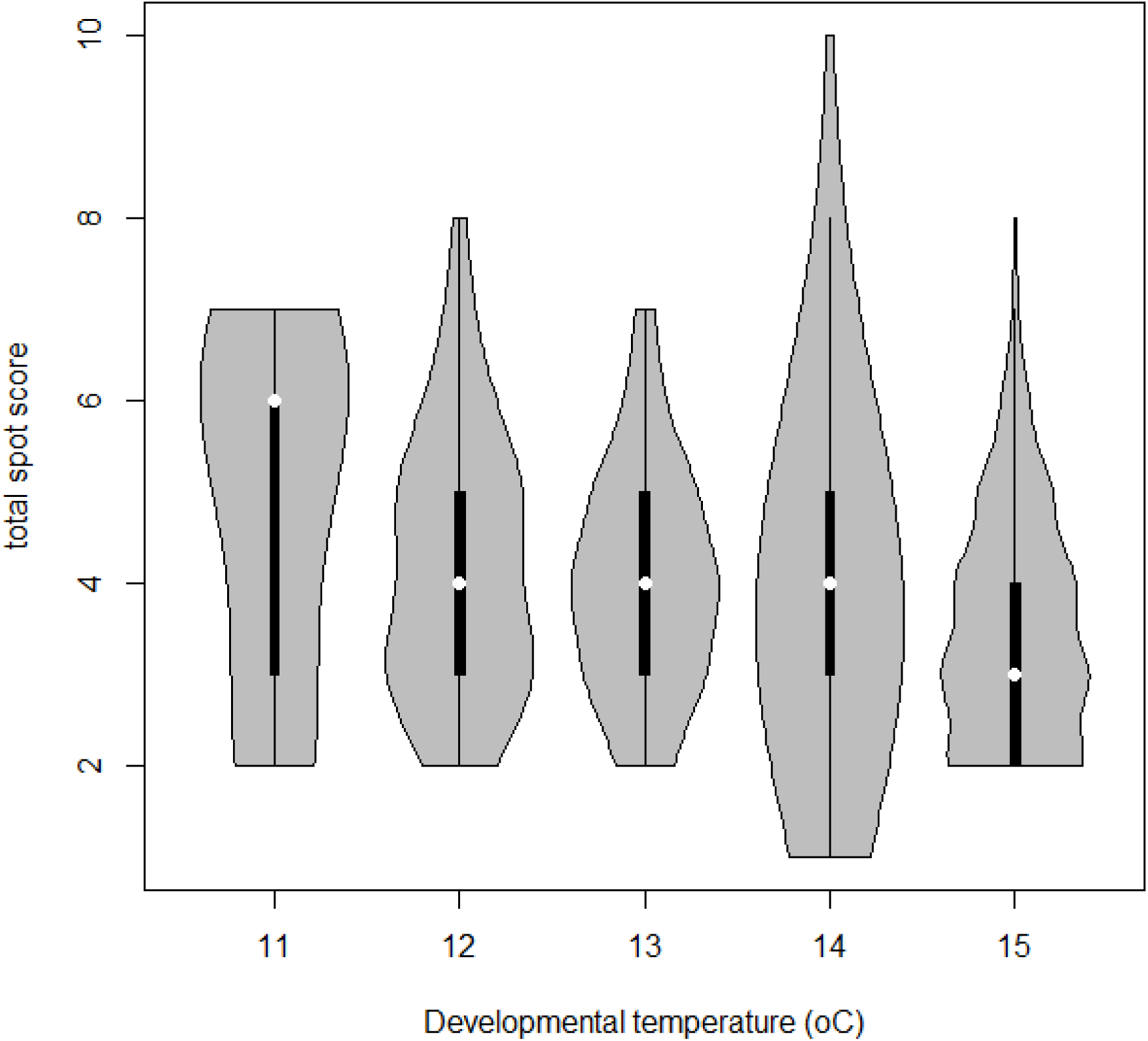
Violin plot of total spot score for all Cornish females collected from Chycoose Farm in 2020, displayed by developmental temperature (rounded to nearest degree C). Note that higher developmental temperature leads to a lower spot score.

**Fig. 3.**
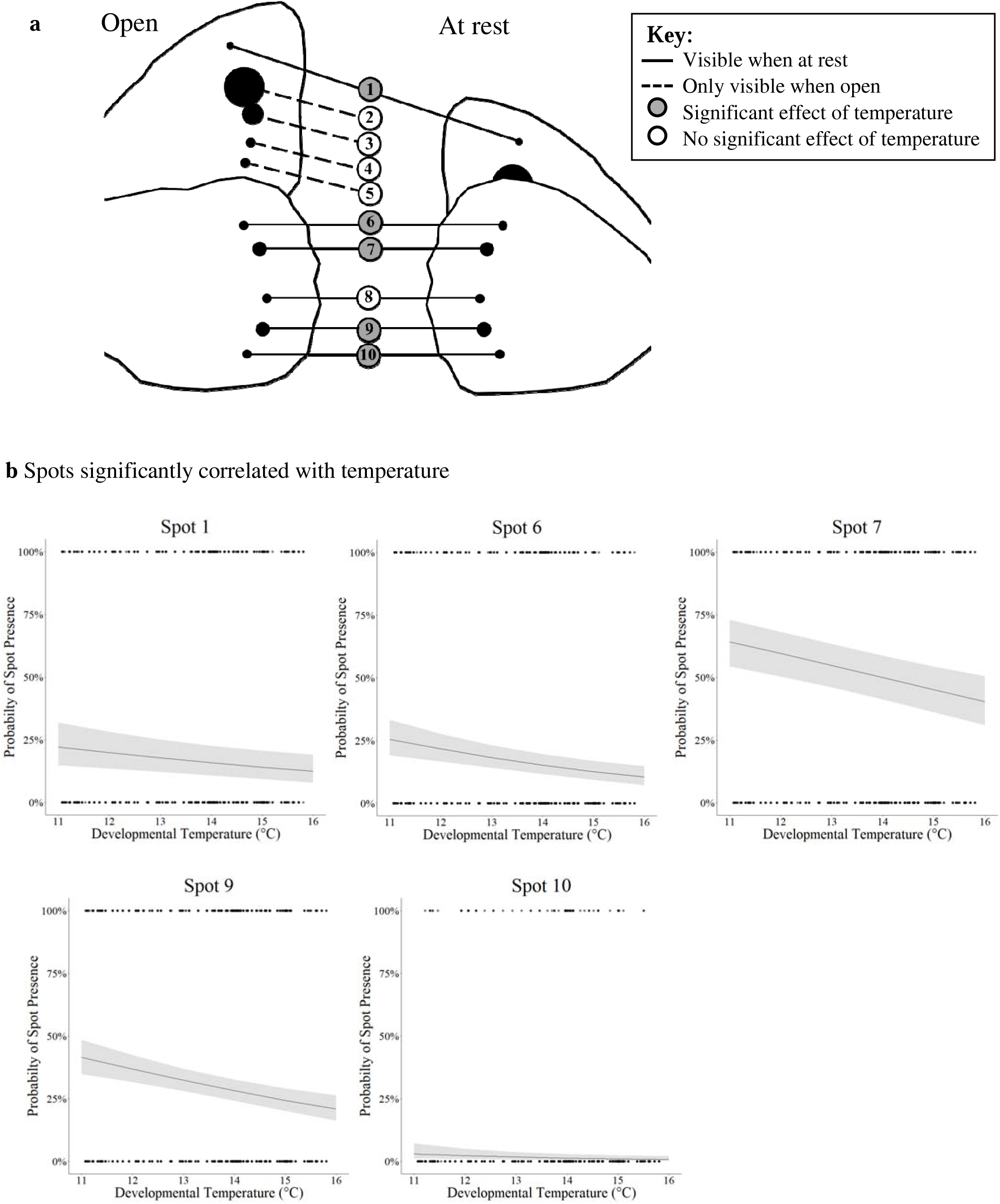
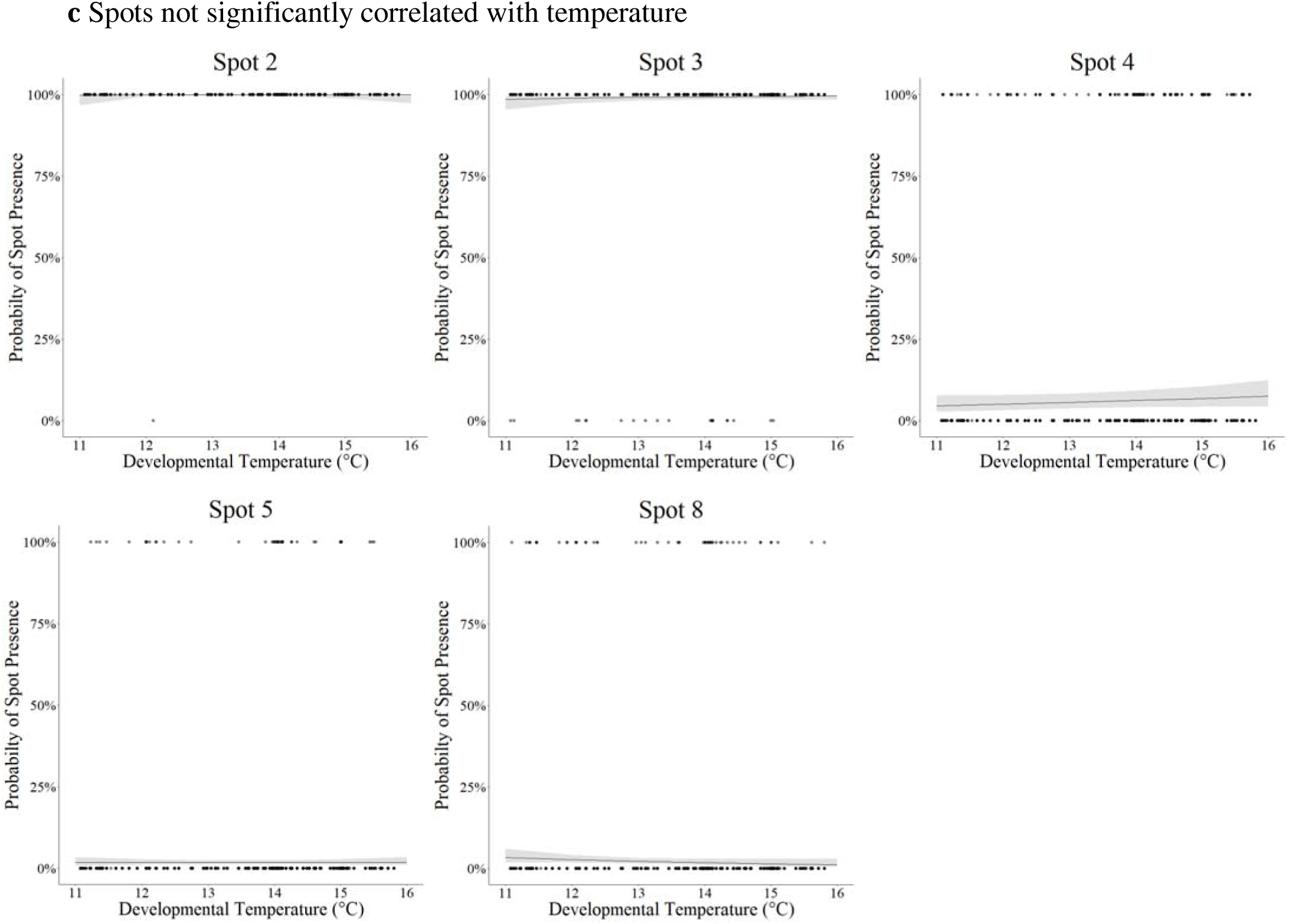
Field temperature determines the probability that wing spots visible in females are present r absent. **a**, Visibility of the ten wing spots (spots 0-10) when the butterfly’s wings are open and when at rest. Showing that spot numbers 2, 3, 4 and 5 are not visible when the butterfly is at rest. **b**, Presence or absence of spot numbers 1, 6, 7, 9 and 10 are significantly affected by temperature (see text). In all five of the spots, the probability of having the individual spot declines as developmental temperature increases. **c**, Presence or absence of spots number 2, 3, 4, 5 and 8 are not significantly affected by temperature. The points show the observed values from the combined Buckingham, Eton and Truro data (n=2,504 for each spot scored), the line shows the fixed effects of the model that spot likelihood is affected by developmental temperature, and the grey shading shows the 95% confidence intervals of the model’s predictions.

### Predicted number of wing spots and intra-seasonal patterns

To predict the likely number of eyespots the predicted developmental temperatures for each of the BMS transect records was calculated using the previously identified lag time and standard deviation from the sliding window analysis. For each of the twenty transects, 5 km resolution HadUK-Grid temperature data was again downloaded from the MET office. Using this developmental temperature, the wing spot total for each butterfly observed was predicted using the model created using the Buckingham, Eton and Truro data. The predicted wing spot totals of the butterflies were then plotted against the day of the year, for the ‘North’, ‘Southwest’ and ‘Southeast’ localities separately. General linear models were subsequently used to statistically test the significance of any change in predicted wing spot totals intra-seasonally.

### Spatial and temporal patterns in predicted wing spot totals

To look at how eyespot variation changes over time and space the annual average predicted wing spot total for each of the transects was calculated; this mean was weighted by the number of *M. jurtina* counted on each of the individual transect walks within that year. This data was then analysed to look for any spatial patterns, firstly examining the eastings and northings of the BMS transects. A general linear model was used to test whether northings of the transect has a significant effect on the annual average predicted wing spot total. This analysis was repeated for the eastings of each transect as well. To examine inter-seasonal patterns in predicted wing spot totals, the Lullington Heath transect (Grid reference Q540020) was used, on the basis that it was walked more frequently than any of the other selected transects (n=668, from 1979 to 2019). The use of a single transect removes the effects of geographical differences, and so instead reflects temporal differences in temperatures. The mean annual average developmental temperatures were calculated; this mean was weighted by the count of *M. jurtina* to reflect the developmental temperatures experienced by most of the population. Data for the average summer temperature was obtained from the MET office, to provide an indication of the general conditions each year. This seasonal summer temperature is the mean temperature value between June and August, extracted at a 5 km resolution for each of the years the transect was walked. GLMs were used to test the relationship between the annual average predicted wing spot total and the seasonal summer temperature; the seasonal summer temperature and annual average developmental temperature; the seasonal summer temperature and the year; and the annual average predicted wing spot total and the year. These models were subsequently plotted as scatter plots, with the model as a regression line. The analysis of the effect of the seasonal summer temperature on annual average predicted wing spot total, and the effect of the seasonal summer temperature on annual average developmental temperatures, was also repeated for the dataset with all the twenty transects, to test if the observed patterns are a generality, or if they exclusive to the Lullington Heath transect. In this analysis, the seasonal summer temperatures will also reflect geographic differences, because multiple transects are used. This seasonal summer temperature data may be a better reflection of the general conditions than eastings or northings, because other factors, such as proximity to the sea and altitude, will also determine the general environmental conditions. This seasonal summer temperature was extracted for each transect location at a 5 km resolution, for each of the years the transect was walked. For the transect at Lindisfarne, 5 km was not a suitable resolution, so temperature data at a 1km resolution was used instead. These general linear models were again subsequently plotted as scatter plots, with the model as a regression line.

## Results

### Field temperatures during pupation correlate with eyespot variation

To look at the effects of temperature on eyespot variation we scored 4,742 butterflies at each of the ten candidate spot positions (Fig. S1), corresponding to the combined scoring of the presence-absence of a spot at 47,420 different wing positions throughout the entire study. Butterflies were collected from three different sites across the entire flight season and over several different seasons (Eton from 1988-1993, n=2,158; Buckingham 1988-1991, n=788 and Truro in 2020, n=1,796 butterflies). First, we confirmed the original observations of Dowdeswell and Ford, that spot numbers in females (n=2,504 butterflies) declined across the season at all three sites (GLM: X^2^_7,2_=14.6, *P*<0.001, Fig. 1a) in so called intra-seasonal variation (Dowdeswell, 1981). Second, we performed a principle-components analysis (PCA) of spot pattern in both sexes combined (Fig. S2). As this showed significant variation in spot pattern between males and females, we ran a new PCA on females only. Principle-component one (accounting for 39% of the variation) in the female only analysis corresponds to variation in the total number of spots (Fig. 1b). Finally, to estimate the local temperature at each site during pupation we used 5km gridded temperature data sets of mean daily temperature for the dates and all three locations studied (see Material and Methods). We performed a GLM to predict the first component of the PCA analysis (spot number) using temperature during a variable period prior to collection date of the butterfly. We then used a moving window analysis to find the period during which temperature had the greatest effect, weighting temperature using a Gaussian weighting. The optimal time window, prior to capture of the butterfly, was chosen as the model with the lowest Akaike information criterion (AIC) value (AIC = 8577.34) to explain variation in spot pattern. Such an analysis for all females shows the most parsimonious model was a sampling window of 35 days, +/- a standard deviation of 8 days (Fig. 1c). This model shows that eyespot variation is driven by the temperature 35 days prior to the capture of the butterfly, at a time consistent with the animal being at the start of pupation (assuming a total of 28 days spent in the pupa (Dowdeswell, 1981; Eeles, 2019) and ∼7 days spent as an adult prior to capture in this study). Here we term this the ‘developmental temperature’, which we predict to be the temperature during early pupation.

To look at the effects of developmental temperature on total spot count across a single season we also analysed the single large and continuous series of Cornish females collected across the 2020 flight season. Plots of total female spot score against developmental temperature for this site (Fig. 2) clearly show that higher developmental temperature leads to a temperature for this site (Fig. 2) clearly show that higher developmental temperature leads to a lower overall spot score. This directly explains Ford and Dowdeswell’s original observation that spotting in females declines over the season, in their so called ‘intra-seasonal’ spot variation (Dowdeswell, 1981).

### Female eyespots visible when the butterfly is at rest are variable

Using this developmental time window (35 days prior to capture), there is a highly significant effect of developmental temperature on PCA component 1, with higher temperatures leading to fewer spots (P=0.001). Analysing spot number totals using the same method, there is also a highly significant effect of developmental temperature on both hind- and fore-wing, and total number of spots (*P<*0.001). Fascinatingly, if the relationship between individual spots and developmental temperature is plotted (Fig. 3), then variation in five of the six spots that are visible in female butterflies at rest (Fig 3a), namely spots 1 (GLM: X^2^_3,1_=7.32, *P=*0.006), 6 (GLM: X^2^_3,1_=17.6, *P<*0.001), 7 (GLM:X^2^_3,1_=19.9, *P<*0.001), 9 (GLM: X^2^_3,1_=21.6, *P<*0.001) and 10 (GLM: X^2^_3,1_=5.18, *P=*0.023) is significantly correlated with temperature (Fig. 3b). With increasing developmental temperature leading to a decrease in the probability that each of these visible spots will be present. Whereas, in contrast, all four of the spots hidden at rest (spots 2 (GLM: X^2^_3,1_=1.00, *p=*0.317), 3 (GLM: X^2^_3,1_=1.42, *p=*0.233), 4 (GLM: X^2^_3,1_=1.81, *p=*0.178) and 5 (GLM: X^2^_3,1_=0.0038, *p=*0.951)) show no significant variation with developmental temperature and neither did spot 8 (GLM: X^2^_3,1_=3.18, *p=*0.07) (Fig. 3c). This data is consistent with the hypothesis that spots visible in the female at rest are under temperature driven developmental control and that this drives the decline in the spottiness of females throughout the season. In contrast, in males only one spot (spot 3) differs significantly with developmental temperature (Fig. S3 and Table S1) and most of the variation observed is driven by location. This suggests that spottiness in males, unlike females, is not under such strong temperature driven control and probably explains why the studies of E.B. Ford and others have only examined variation in females (Dowdeswell, 1981; Dowdeswell et al., 1949; Dowdeswell and Ford, 1955; Dowdeswell et al., 1957; Dowdeswell et al., 1960; Dowdeswell and McWhirter, 1967), as these are indeed more variable in the field.

### Wing length scales with the size of the large forewing spot 2/3

Monthly temperatures have recently been correlated with wing length (or ‘body size’) in several species of UK butterflies (Davies, 2019). Moreover, other studies of eyespot plasticity in different butterflies also recognise that wing length can affect overall eyespot size (total area) and therefore in many recent studies eyespot area is often expressed relative to wing length (Brakefield et al., 1996). This observation is particularly relevant to the large compound eyespot, here termed spot 2/3 (termed ‘compound’ as it is invariably a merger of spots at positions 2 and 3, see Material and Methods), on the forewing of both sexes. Because this large spot is never absent (it was present in all specimens of both sexes in the current study) we examined the effect of wing length on the height, width and estimated total area of spot 2/3. We find that forewing length does indeed decrease across the flight season in the large cohort of female butterflies collected from a single site in Cornwall in 2020 (see Supplemental Fig. S4). However, critically, wing length variation is best explained by the day of the year rather than developmental temperature which accounts for the variation in the presence/absence of the remaining eight smaller spots. Thus, a linear model predicting wing length as a function of both developmental temperature and day of the year shows a significant effect of day of year (wing length decreases through the season, P<0.001; Figure S4) but no effect of developmental temperature (P=-0.254). The partial r^2^ (explanatory power) of day of year (0.001, Table 1 below) is therefore considerably higher than that of developmental temperature (0.019, Table 1). Critically, this suggests that wing length is controlled independently of spot presence/absence (see also forewing size data displayed in Supplemental, Figs. S5-S6 and statistical analysis reported in Tables S2-S3). This further analysis confirms that total spot count (the central character scored by E.B. Ford and others) is correlated with developmental temperature whereas wing length and the corresponding size of the large omni-present spot 2/3 is correlated with day of the year and is therefore likely to be controlled independently.

**Table 1.** Day of the year is a better predictor of wing length than developmental temperature.

| | Coefficient | Std. error | t value | P value | partial $r^2$ |
| --- | --- | --- | --- | --- | --- |
| Intercept | 27.892782 | 0.459468 | 60.707 | < 2e-16 *** |  |
| Developmental temperature | 0.094210 | 0.082596 | 1.141 | 0.254 | 0.001 |
| Day of year | -0.022635 | 0.005136 | -4.407 | 1.16e-05 *** | 0.019 |

### Wing damage and fore- and hindwing spotting

Eyespots on butterfly wing margins are thought to be attractive to predators and therefore to deflect attacks onto the wings and away from the body (Ho et al., 2016). However, given that spottiness decreases across the season we wanted to examine the likely effects of this decline on predation (wing damage). To understand the relationship between eyespots and wing damage over the season we looked at the Eton samples in more detail. We recorded the total number of spots per wing (fore- and hindwing) and the presence or absence of damage on each wing, scored as symmetrical (inferring an attack when wings are closed on the ground) or asymmetrical (inferring at attack on a single wing in the air). Damage in all recorded categories (forewing, hindwing, symmetrical, asymmetrical and total damage) increased through the season (*P*<0.0001 in all cases). Therefore, to look at the role of eyespot number on wing damage independently of time we carried out a binomial linear regression with day of the year as a covariate. This analysis (Fig. 4) shows that more hindwing spots lead to less asymmetrical damage in both males and females, suggesting they may indeed confuse predators in flight. Second, more forewing spots led to less symmetrical damage in males but more asymmetrical damage in females. This suggests that males may use forewing spots to flash at predators when at rest, whereas more forewing spots in females may make them more obvious to predators in flight. Given the strong decline in hindwing spots in both males (*P*<0.0001) and females (*P*<0.0001) across the season, this suggests that reduced hindwing spotting at the end of the season would be expected to increase aerial predation of both sexes.

### Phenology of butterfly flight across the UK

Given that spottiness in both sexes declines with increasing developmental temperature in the field, we wanted to examine if the Meadow Brown can effectively compensate for such changes in temperature by altering its flight phenology across its range. We therefore examined flight period data from the Butterfly Monitoring Scheme from 20 transects (Table S2), selected from across the United Kingdom (Fig. 5a, Table S2) and each sampled for 40 years continuously. The phenology of the flight season differed between the localities sampled but only towards the end of the season (Fig. 5b). The timing of peak population sizes was therefore consistent between sites (the 50% quantile varies from day 145 to 152 between the localities; Table S2), as was the start of the flight season (2.5% quantile varies from day 116 to 119 between the localities; Supplemental Table S2), consistent with previous studies (Brakefield, 1987). But the timing of the end of the flight season, however, varies significantly between locations, being earlier in the Northern transects (97.5% quantile is day 175 for the North transects, compared to day 191 and day 192 for the Southeast and Southwest, respectively). This means that the length of the flight season is shorter in the more northerly transects; the 95% range for the North transects is 56 days, compared to 74 days and 75 days for the Southwest and Southeast respectively. Finally, we used this data to predict female spot number variation in the 20 sampled transects. The predicted wing spot total declined significantly with day of the year in the North (GLM: X^2^_3,1_=665.8, *P*<0.001; Figure 4c), Southwest (GLM: X^2^_3,1_=2701.6, *P*<0.001; Figure 5c) and Southeast (GLM: X^2^_3,1_=2582.8, *P*<0.001; Figure 5c) transects. Intra-seasonal shifts for lower wing spot totals are therefore predicted across the country, regardless of location.

**Fig. 4.**
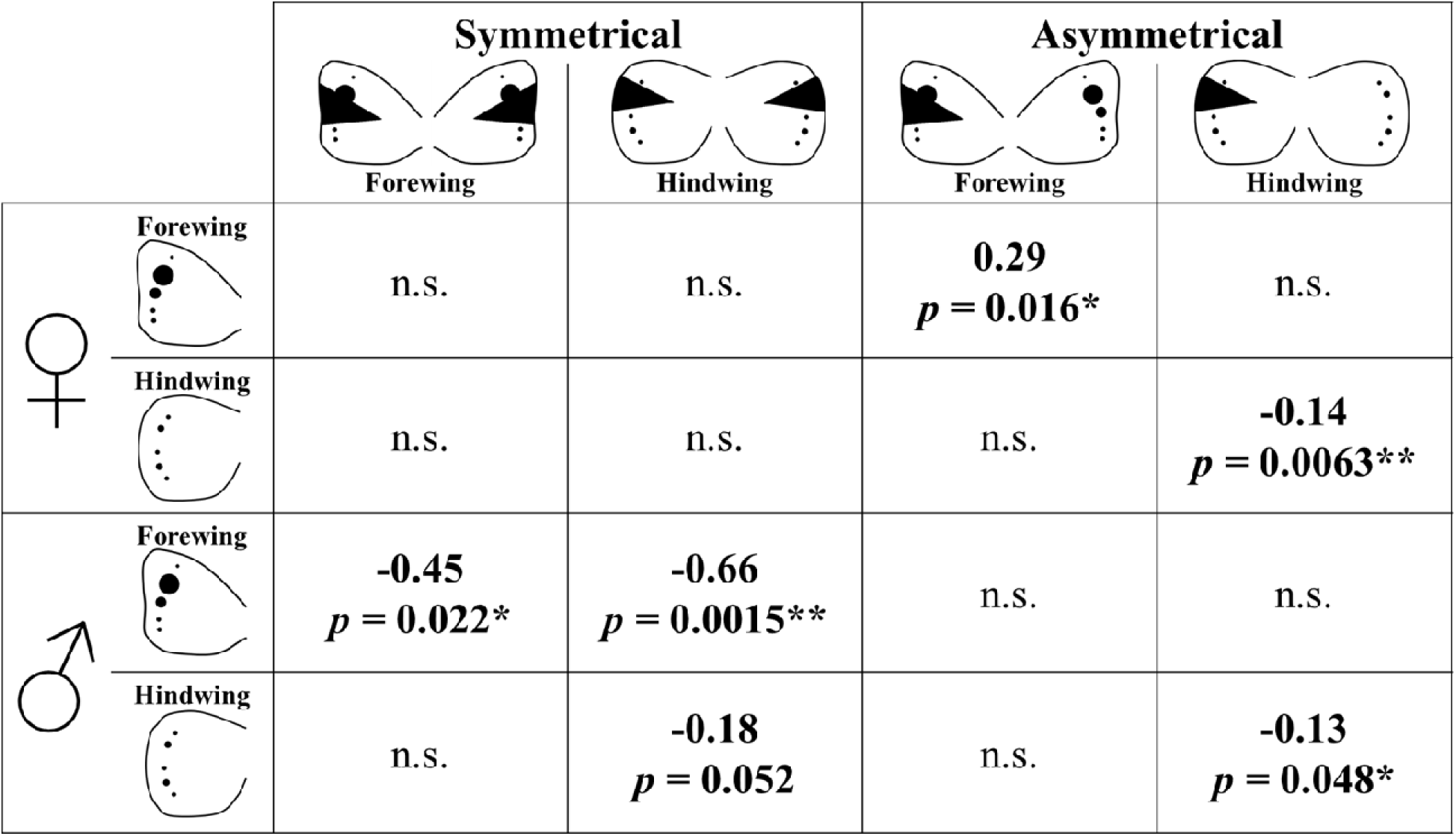
Relationship between spot number and wing damage for fore- and hindwing in males and females. Results of a binomial regression analysis showing the relationships (negative or positive) between damage (symmetrical or asymmetrical) and spot number on each wing. See text for discussion.

**Fig. 5.**
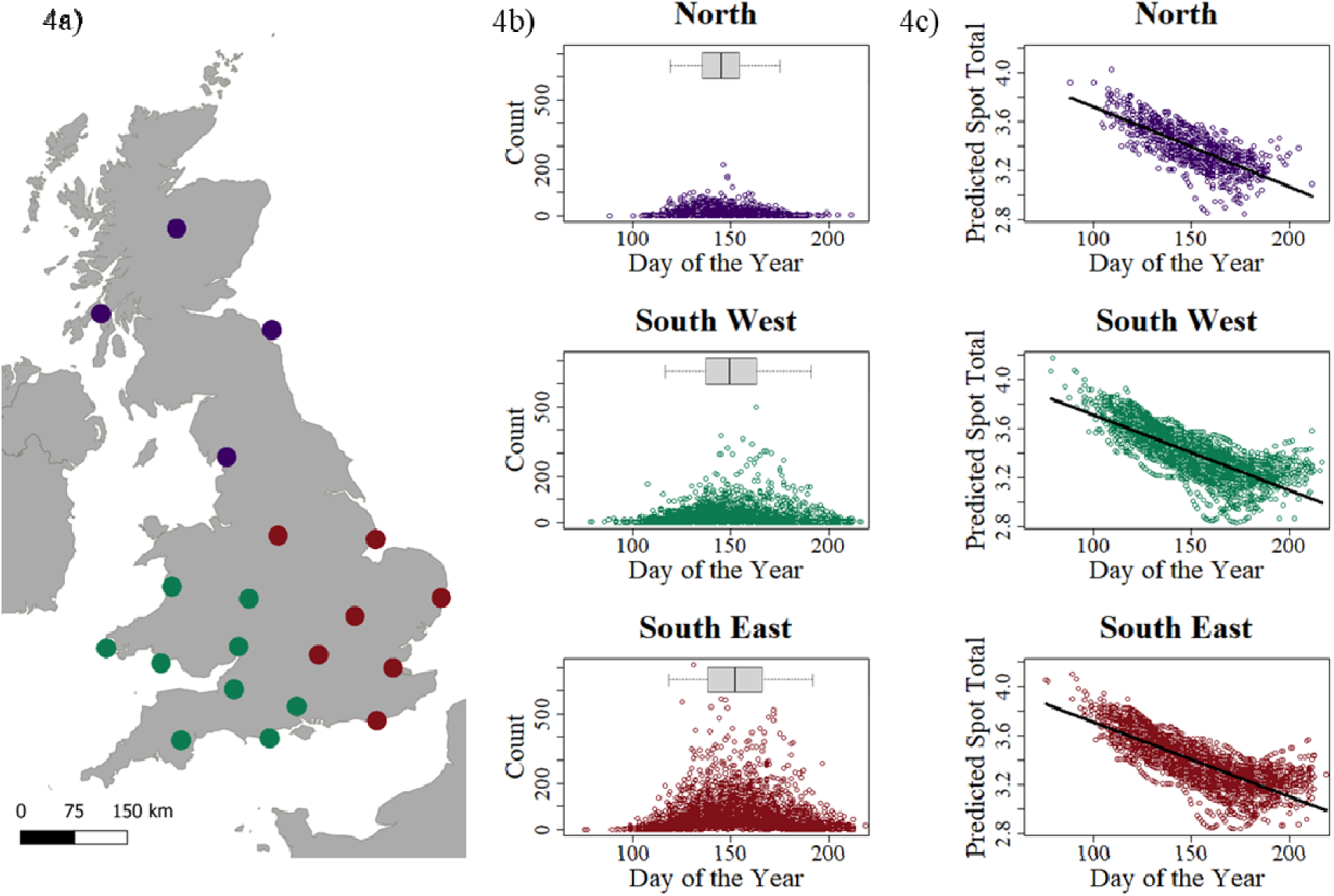
Phenology of the Meadow Brown flight period across the UK and the corresponding decline in female spottiness predicted intra-seasonally. **a**, Location of the twenty selected BMS transects, with the colours representing whether they were classified as ‘North’ (purple), ‘South West’ (green) or ‘South East’ (red). **b**, Number of Meadow Browns observed in each transect walk, against the day of the year the transect was walked, for each of the localities. The day of the year is the number of days since the start of the flight season on 1 March. The box plots above each plot (in panel **b)** show the quantiles of the day of the year data. The central line is the 50% quantile, the edges of the box are the 25% and 75% quantile, and the whiskers are the 2.5% and 97.5% quantiles. **c**, Predicted spot totals change with day of the year at each of the localities. The predicted spot total declines intra-seasonally in all three localities.

### Warmer summers predict fewer female spots

To predict how increasing summer temperatures might decrease female spottiness we looked at a single BMS transect from Lullington Heath, in the South of England. The seasonal summer temperature at this site is predicted to have a significant effect on annual average predicted wing spot total (GLM: X^2^_3,1_=24.5, *P*<0.001; Fig. 6a) with female spottiness decreasing with increasing temperature. The seasonal summer temperature also has a significant effect on annual average developmental temperature with developmental temperatures increasing as summer temperatures increase (GLM: X^2^_3,1_=25.2, *P*<0.001; Fig. 6b). However, unexpectedly this is not a simple one-to-one relationship (compare the red and blue lines in Fig. 6b) and therefore seasonal temperature alone is a poor predictor of spottiness. Seasonal summer temperatures have increased significantly over the 40-year span that the transect has been surveyed (GLM: X^2^_3,1_=12.8, *P*<0.001; Fig. 5c). Finally, therefore, year also has a significant effect on the annual average predicted wing spot total (GLM: X^2^_3,1_=7.40, *P*=0.006; Fig. 6d) with the annual average predicted wing spot total predicted to decrease over time. Analysis of all twenty of the selected BMS transects also found there is a significant effect of seasonal summer temperatures on the annual average predicted wing spot total (GLM: X^2^_3,1_=22.8, *P*<0.001; Fig. 7a). However, the relationship is unexpectedly positive; as the seasonal summer temperature increases, the predicted number of wing spots increases. There is a significant effect of seasonal summer temperatures on developmental temperatures across all the transects (GLM: X^2^_3,1_=48.6, *P*<0.001; Fig. 7b). Taken together this data shows that seasonal summer temperature alone is a poor predictor of spot variation, suggesting that care needs to be taken when selecting the correct variable to predict the effects of climate warming on phenotypic change.

**Fig. 6.**
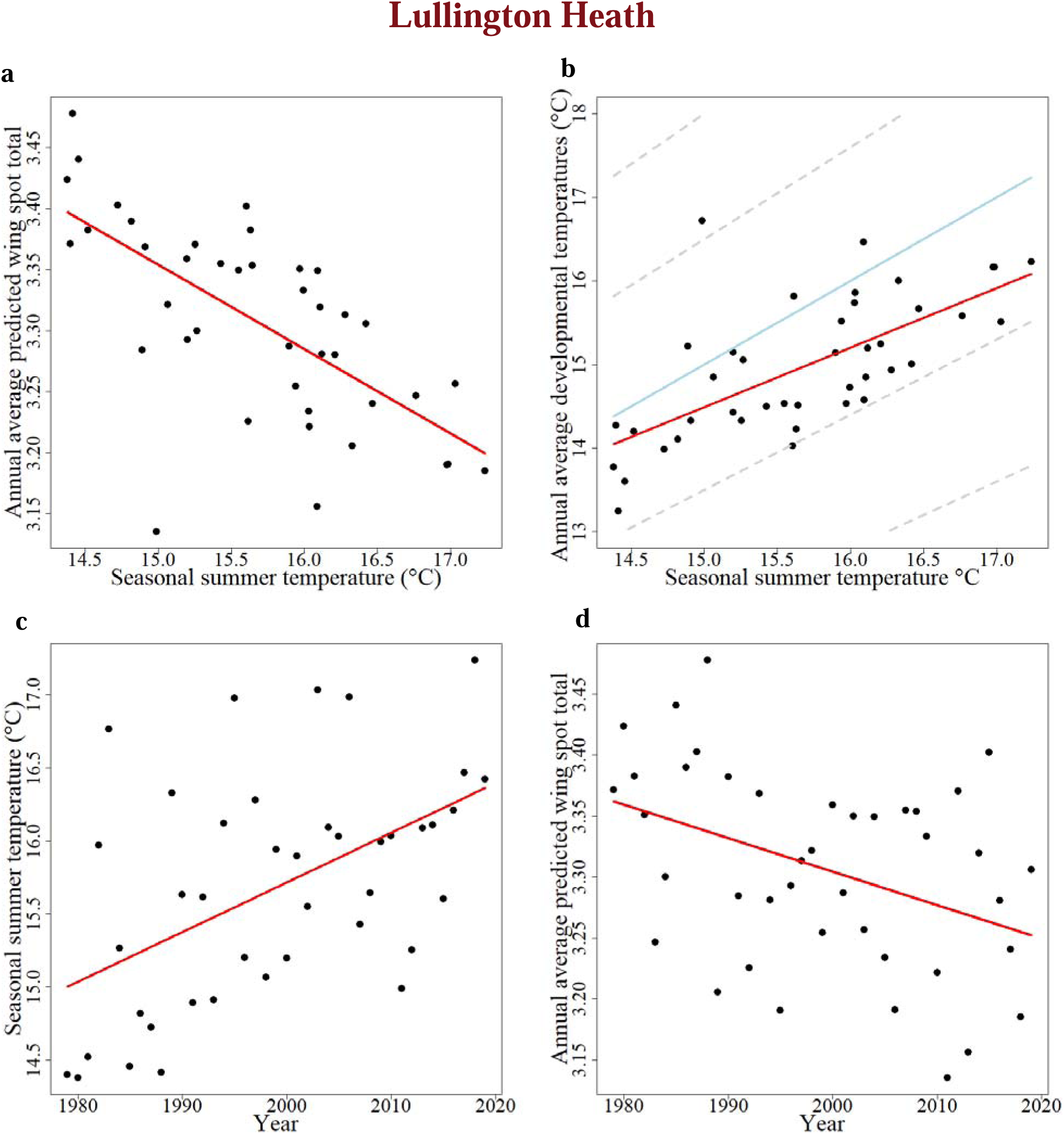
Temperature data over multiple years for a single site in Southern England (Lullington Heath) predicts a long-term decline in female spottiness. **a**, Average predicted wing spot total declines as seasonal summer temperatures increase. Each point shows a different year the transect was walked, and the red regression line shows the model. **b**, Annual average developmental temperature increases as seasonal summer temperatures increase. The red line shows the slope of the linear model for the relationship between the developmental and summer temperatures. Note, the slope of this line differs to that of a one-to-one ratio shown by the blue line (see text). The grey dashed lines show a 1 to 1.4, a 1 to 1.2, a 1 to 0.8, and a 1 to 0.6 ratio from top to bottom respectively. **c**, Seasonal summer temperatures have increased significantly over the 40 years the Lullington Heath transect has been walked. **d**, The average predicted wing spot total declines over time with warmer summers.

**Fig. 7.**
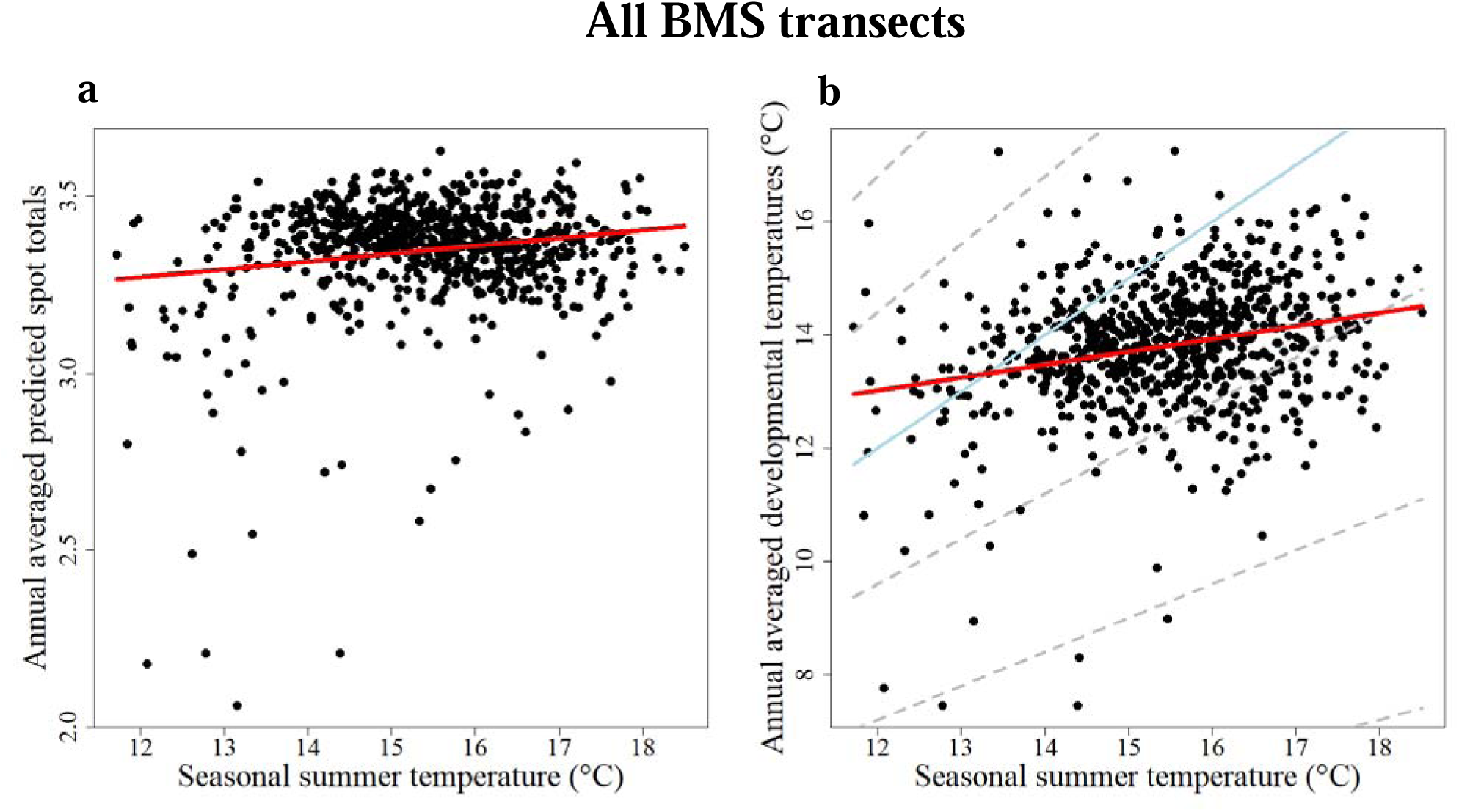
Seasonal temperature is a poor predictor of spot total and developmental temperature. Data from all twenty BMS transects is combined to examine how good a predictor seasonal summer temperature is of spot variation. **a**, Annual averaged predicted number of spots increases as seasonal summer temperatures increase. **b**, Annual averaged developmental temperature is correlated positively to the seasonal summer temperature. The red line shows the slope of the linear model for the relationship between the developmental and summer temperatures. The slope of this line differs to that of a one-to-one ratio (shown by the blue line). The grey dashed lines show a 1 to 1.4, a 1 to 1.2, a 1 to 0.8, a 1 to 0.6 and a 1 to 0.4 ratio from top to bottom respectively. The points each correspond to a single transect in a single year. The equation of this model is: y = 10.28+0.22x, where y is the developmental temperature and x is the seasonal summer temperature

## Discussion

Our results show that variation in the number of spots in Meadow Brown butterflies is related to field temperature, both within and between seasons. This finding stands in contrast to the previous 80 years of work by others in which authors have suggested that variation is associated with a genetic polymorphism and generated by the differential survival or development of high and low spot ‘morphs’ (Baxter et al., 2017; Beaufoy et al., 1970; Bengston, 1978; Brakefield and van Noordwijk, 1985; Conradt et al., 2000; Creed et al., 1964; Dowdeswell et al., 1949; Dowdeswell and Ford, 1955; Dowdeswell et al., 1957; Dowdeswell et al., 1960; Dowdeswell and McWhirter, 1967; Frazer and Willcox, 1975; Grill et al., 2007; McWhirter, 1969; Scali, 1972). Limited laboratory experiments varying temperature during pupation have previously failed to show a role of temperature in spotting (Brakefield and van Noordwijk, 1985). However, the role of field temperature inferred here now helps explain why drought (macroclimate) or changes in grazing (microclimate) might affect total spotting and may also explain why spot number can appear to vary on either side of single hedgerow (Creed et al., 1959), perhaps now best explained by microclimate rather than strictly by population genetics. Our finding that the critical time window for determining spot number is 35 days prior to capture (Fig. 1c) is consistent with the butterfly spending 28 days in the pupa(Dowdeswell, 1981; Eeles, 2019) and ∼7 days on the wing prior to capture, placing this time window at the anticipated point of early pupation, which is the critical developmental period for butterfly wing pattern determination (Beldade and Brakefield, 2002; Beldade et al., 2002). This observation is also consistent with a recent comparative study of eyespot variation in other satyrid butterflies, where temperature has also been shown to drive eyespot size variation via the action of the insect hormone 20-hydroxyecdysone (Bhardwaj et al., 2020). When we look at the effect of temperature on each individual female spot-position we find that only those spots that are visible, when the butterfly is at rest, vary in their presence/absence with developmental temperature and the spots that are hidden do not vary significantly (Fig. 2b versus c). This is consistent with the hypothesis, derived from the tropical butterfly Bicyclus anynana (Lyytinen et al., 2004), that predation maintains eyespot plasticity. It is also consistent with the idea that ‘covered’ eyespots, which are not continuously visible to predators, are less likely to be under positive selection (Chan et al., 2021).

Paul Brakefield (Brakefield and Shreeve, 1992) stated that if temperature alone drives spot-formation then the spots themselves will then be of no evolutionary significance. This historical argument, however, overlooked the more recently recognised adaptive role of environmental plasticity (Ghalambor et al., 2007). We therefore used wing damage and spot number to examine if seasonal plasticity affects selection via predation (as measured by wing damage). The effect of temperature driven phenotypic plasticity is stronger in females versus males, however, declines in hindwing spottiness across the season are clear in *both* sexes (remembering that Ford only examined females). Examination of symmetrical and asymmetrical wing damage in the Eton dataset (controlled for day of the year) shows that more hindwing spotting leads to less asymmetrical damage in *both* sexes suggesting spots may indeed confuse aerial predators. Whereas more forewing spots produce less symmetrical damage in males but more asymmetrical damage in females, intriguingly suggesting that they may have *opposite* effects in different sexes. A similar relationship between forewing spotting and survival has been noted in *B. anynana* where predation experiments have confirmed that female butterflies with fewer forewing spots live longer and lay more eggs (Chan et al., 2021). Given that the number of hindwing spots in both sexes of *M. jurtina* declines strongly across the season, this suggests that any role in predator deflection will also be decreased over time. The finding that intra-seasonal variation is driven by field temperature therefore suggests that phenotypic plasticity may set up a ‘trade-off’ between the benefits of predator avoidance *early* in the season and the costs in terms of developmental time necessary to mature visible eyespots later in the season. Finally, we modelled the effects of increasing summer temperatures both in a single site at Lullington Heath (Fig. 6) and across a series of UK BMS transects combined (Fig. 7). This analysis predicts that warmer summer temperatures will indeed lead to less spotty females (both withing and between years) but strikingly they also show that seasonal summer temperatures are a poor predictor of developmental temperature and are therefore not likely to predict spot variation on their own. This demonstration of temperature driven phenotypic plasticity therefore not only stands in contrast to previous work predicting that spot variation is explained by population-genetics, but it also suggests that our warming climate will drive further loss of butterfly spottiness year on year across the UK.

## Acknowledgements

We thank Philip Lewis for setting all specimens.

## Data availability

Data and R codes are available at https://github.com/eyespots-in-meadow-brown

